# *Escherichia coli* cells are primed for survival before lethal antibiotic stress

**DOI:** 10.1101/2022.11.11.516214

**Authors:** Tahmina Hossain, Abhyudai Singh, Nicholas C. Butzin

## Abstract

Non-genetic factors can cause significant fluctuations in gene expression levels. Regardless of growing in a stable environment, this fluctuation leads to cell-to-cell variability in an isogenic population. This phenotypic heterogeneity allows a tiny subset of bacterial cells in a population, referred to as persister cells, to tolerate long-term lethal antibiotic effects by entering into a non-dividing, metabolically altered state. One fundamental question is whether this heterogeneous persister population is due to a pre-existing genetic mutation or a result of a transiently-primed reversible cell state. To explore this, we tested clonal populations starting from a single cell using a modified Luria–Delbrück fluctuation test. Through we kept the conditions the same, the diversity in persistence level among clones was relatively consistent: varying from ∼60-100 and ∼40-70 fold for ampicillin (Amp) and apramycin (Apr), respectively. Then we divided and diluted each clone to observe whether the same clone had comparable persister levels for more than one generation. Replicates had similar persister levels even when clones were divided, diluted by 1:20, and allowed to grow for ∼5 generations. This result explicitly shows a cellular memory passed on for generations and eventually lost when cells are diluted to 1:100 and regrown (>7 generations). Our result demonstrates 1) the existence of a small population prepared for stress (“primed cells”) resulting in higher persister numbers, 2) the primed memory state is reproducible and transient, passed down for generations but eventually lost, and 3) a heterogeneous persister population is a result of a transiently-primed reversible cell state and not due to a pre-existing genetic mutation.

## Introduction

Clonal populations often exhibit phenotypic heterogeneity, leading to specific physiological effects that distinguish some cells from others^1-3^. Variability can slightly reduce fitness in a common environment but, in return, maximize it during environmental perturbations^2, 4^. For example, bacterial persister cells, a phenotypic variant, can endure prolonged lethal antibiotic treatment by entering a metabolically altered state^5^. This small subpopulation can reestablish infection after treatment and necessitate repeated long-term antibiotic therapy. They also have a high mutation rate that increases the likelihood of evolving antibiotic resistance, a pressing public health concern. A recent study provides evidence that among infectious diseases, antibiotic resistance may be the leading cause of death worldwide (more than HIV or malaria)^6^. Global death projections are now estimated at 100M/year by 2050^7, 8^ unless new technologies are developed to combat them, and we have a better understanding of persister survival. Here, we specifically focus on how heterogeneity in a population drives persister numbers.

Noise or fluctuation in gene expression levels drives phenotypic variation that could be an inherent survival strategy of a clonal bacterial population^2, 9^. Studies reported that stress-response genes are generally more variable than housekeeping genes^10, 11^. But it is uncertain whether this variability is controlled or an effect of inevitable stochastic fluctuations in gene expression^12^. We formalize this notion and hypothesize that specific cells are prepared for stress through a transiently inherited cell state.

In this work, we take advantage of the Luria & Delbrück fluctuation test (FT), which was recently used to identify genes related to cancer persistence against cancer drugs^13^. Cancer persisters are similar in phenotype to bacterial persistence but biochemically unrelated. We used a similar approach to probe bacterial persistence. The FT was first pioneered ∼80 years ago when Luria & Delbrück demonstrated that genetic mutations arise randomly in the absence of selection but not in response to the selection^14^. They were studying phage (bacterial virus) infections. During that time, it was debated whether mutations leading to resistance were directly induced by the virus (Lamarckian theory) or if they developed randomly in the population before viral infection (Darwinian theory). They designed an elegant experiment where single cells were isolated and grown into clones and then infected by a phage (Fig. 1a). The number of resistant bacteria was counted across clones. Suppose mutations are virus-induced (i.e., no heritable genetic component to resistance), where each cell has a small and independent probability of acquiring resistance. In this case, clone-to-clone fluctuations in the number of resistant cells should follow Poisson statistics (a memoryless process). In contrast, if mutations occur randomly before viral exposure (spontaneous mutation), the quantity of surviving bacteria will vary considerably across clones depending on when the mutation arose in the clone expansion. The data clearly showed a non-Poissonian skewed distribution for the number of resistant bacteria, validating the Darwinian theory of evolution^14^ (Fig. 1a). This work led to a Nobel Prize, and the FT remains the most commonly used method to measure mutation rates in microbes^15^. While Luria-Delbrück focused on irreversible genetic alterations driving phage resistance, here we use the FT to elucidate transient cell states that originate via reversible non-genetic mechanisms^13, 16-18^. Notably, this generalized FT can be applied when single cells reversibly switch between drug-sensitive and drug-tolerant states even before treatment. Clone-to-clone fluctuations can be exploited to quantify these switching rates rigorously^19^. This approach has been key in deciphering drug-tolerant cancer cells that arise stochastically even before drug exposure. Thus cancer cells have a bet-hedging mechanism to survive sudden hostile extracellular environmental changes^13, 16, 17^. Here, we are exploring ‘primed cells’ (cells prepared for stress and arise by a rare, transiently inherited cell state) present before the treatment and prolonging survival once treatment begins (Fig. 1b-e).

**Fig. 1.**
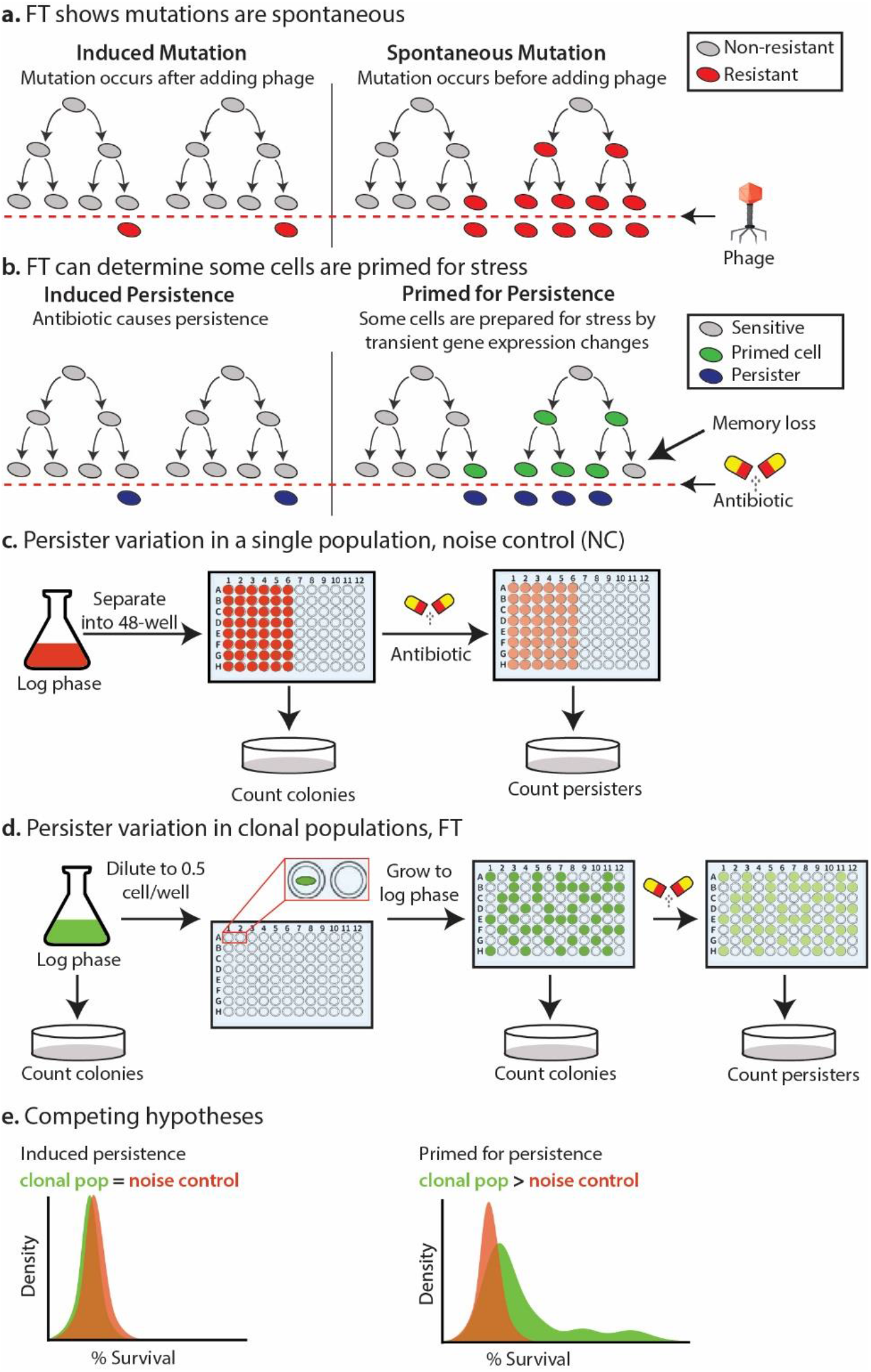
**a**. The Luria-Delbruck fluctuation test (FT). Each clone starts from a single cell and is then infected by a phage (virus). If resistance mutations are virus-induced, the number of resistant cells would follow a Poisson distribution across clones. In contrast, if mutant cells arise spontaneously prior to viral exposure, there will be considerable clone-to-clone fluctuations in the number of surviving cells, including mutations that happen early in the lineage expansion causing many cells to be resistant. **b**. FT to determine if some cells are primed for stress. This design follows the same experimental setup as a., but it uses antibiotic stress and measures variation in persister cells (cells in an altered cellular activity state by having transient gene expression change) across the clones. If persisters are antibiotic-induced, then the number of persister cells would follow a Poisson distribution across clones. In contrast, if there are primed cells with a unique transient gene expression profile (caused by non-genetic factors) that allows them to prepare for stress before antibiotic exposure, there will be considerable clone-to-clone fluctuations in the number of surviving cells. Transient gene expression changes can be heritable and happen early in the lineage expansion, causing many cells to become primed for stress with a transient memory. **c**. Persister variation check in a single population, referred to as NC: Noise control. Cultures were grown to mid-log phase, ∼1E+8 CFU/ml, separated into 48 wells, treated with an antibiotic, 0.1 mg/mL Amp or Apr, and plated before and after antibiotic treatment to get % Survival. **d**. Persister variation check in clonal populations started from a single cell, referred to as FT. Cultures were diluted to 0.5 cell/well and treated similarly to a. **e**. A model for competing hypotheses; *Induced persistence:* persisters are induced due to stress, and no difference will be observed in persister variation between clonal populations and noise control. *Primed persistence:* some cells are primed prior to stress; thus, some clonal populations will have more primed cells than others, and the persister variation will be higher in the clonal populations than in the Noise control (NC).

In this study, we applied the FT to indicate some cells are prepared for stress and have an inherent transient memory. We applied this approach in conjunction with mathematical modeling to elucidate stochastic phenotype-switching in response to antibiotic stress, an inherent survival strategy that gives flexibility to a clonal population. Our work demonstrates how heritable transient cell state changes can lead to variation in persister numbers. Exploring this phenomenon can shed light on how bacteria endure stress, a key question in persister research^20^.

## Result

### Primed cells are prepared for stress before antibiotic treatment

Bacterial populations are heterogeneous, even in log growth phase in a well-controlled lab environment, and especially in natural systems^21, 22^. The quantification of persister numbers often have huge variations with large error bars (SD or SEM), sometimes with 100s of fold differences^23-25^. We used defined media (MMB+^26, 27^), which contains only chemically known components, to minimize variations between experiments. We optimized and standardized our experimental design, and reduced the internal error rate to below 2-fold. Much of this error comes from the compounding imprecisions from multiple pipetting (all pipets have an error rate, and these experiments require serial dilutions). Despite reducing the error rate, we noticed that there would occasionally be huge outliers (up to 100-fold higher or lower than the average).

We determined the typical persister variation in an *E. coli* population (Noise control: NC) is higher than our internal error. We grew a culture to mid-log phase and then divided it into 48 wells. We then treated it with ampicillin (Amp) or apramycin (Apr) for 3 h, followed by a persister assay (Fig. 1c). This resulted in a variation of ∼6-fold for either antibiotic. These results do not explain the 100-fold changes we occasionally observed. Unable to explain the variation at the population level, we wondered if a “memory” was passed down over several generations and skewed our data. We set out to test for variations from a single cell using a FT^13, 16, 17^. Cultures were diluted to about 0.5 cell/well (Limiting Dilution assay^28, 29^) in a 96-well microplate; on average, 48 wells had growth, and 48 wells did not (Fig. 1d). Cells were then grown to an OD of 0.4-0.6 (log phase) and treated with Amp for 3 h. Most (but not all wells) reach the desired OD range simultaneously. We use two 96-well plates and pick 48 clones within the OD range to deal with this. After treatment, the cells were plated on Petri dishes, grown, and colonies were counted to get CFU/mL. We first tested lethal Amp dosages for 3 h, and the persister range was vast, ∼60-100-fold. We tested this several times (Fig. 2a; 8 separate experiments with ∼48 clones/experiment), and there is consistently an extensive range of persister variation among the clones. We wondered if this was specific to Amp, so we tested another class of antibiotic, Apr. Amp targets the cell wall^30^, while Apr targets the 30S ribosomal subunit^31^. Again, the persister range was vast, ∼40-70-fold range with Apr (Fig. 2a). We hypothesize that the extensive range in persistence is due to either (1) mutation or (2) some cells are prepared for stress (“primed cells”), and these primed cells exhibited specific characteristics which allow them to prepare prior to the stress. To assess if a mutation causes this extensive persister range, we tested for antibiotic resistance by streaking clones on antibiotic plates, and no resistant colonies grew (Fig. S1a). To further test for mutations, we diluted the high persister clone (Hp clone) into 0.5 cell/well and repeated the fluctuation test. If a high persister clone was mutated, the average persister percentage would increase, and the range would decrease. However, they had the same average persister percentage and similar range (∼60-100 fold) in both fluctuation experiments (Fig. 2b *i*). If the cells developed resistance, their minimum inhibitory concentration (MIC, the minimal antibiotic required to hinder growth) would change. But it remained the same (Fig. 2b *ii*). These results undoubtedly rule out mutations as the cause and led us to test hypothesis 2, that some cells are primed prior to stress.

**Fig. 2.**
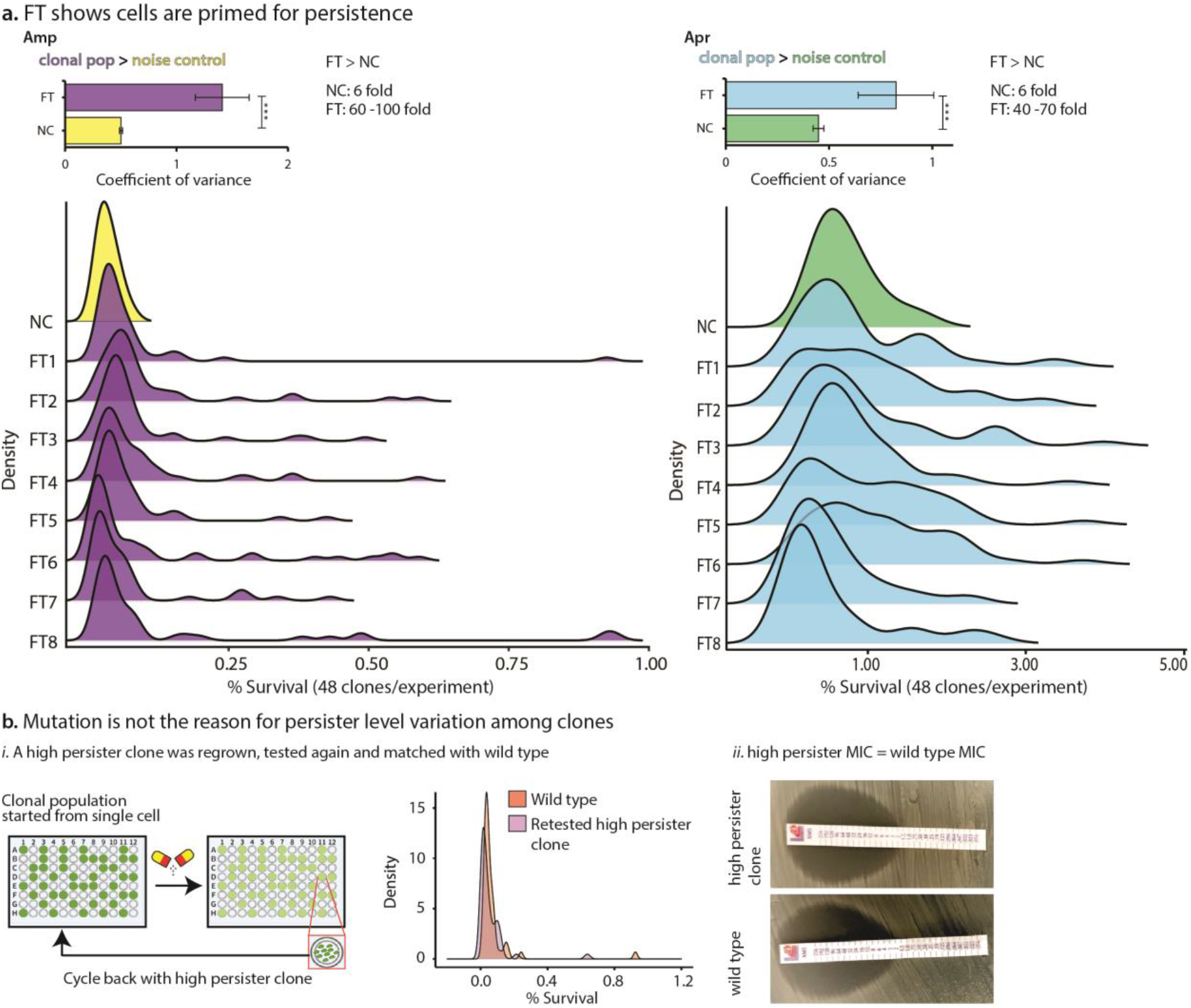
Some cells are prepared and primed for stress. **a**. Results support primed persistence hypothesis: Population-level variation for Amp and Apr: ∼6 fold (experimental setup described in Fig. 1d). FT from single cell level (experimental setup described in Fig. 1e): treatment with Amp or Apr shows 60 to 100-fold and 40 to 70-fold variation, respectively. **b**. Mutations do not cause persister level variation among the clones. ***i***. FT with wild-type clonal populations and FT with clonal populations started from a high persister clone have a similar average persister level and persister range among 48 clones. ***ii***. MIC test showing no change in MIC level in high persister clones. \*\*\**P<0*.*001*.

**Fig. 3.**
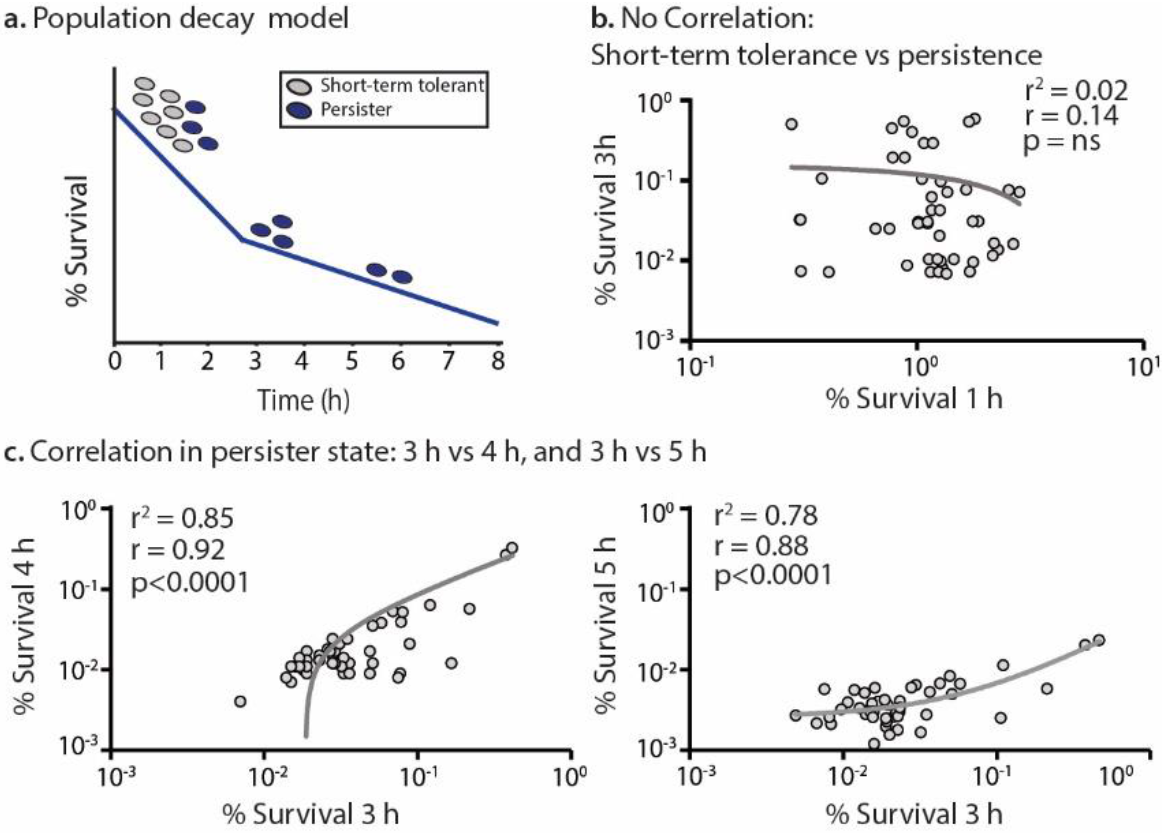
Cells are primed for persistence, not for short-term tolerance. *i*. A simplified model of population decay indicates a biphasic death curve where susceptible cells or short-term tolerant cells die quicker than persisters. ***ii***. Compare each clone’s short-term tolerance levels (1 h Amp) to their persister level (3 h Amp). ***iii***. Compare each clone’s persister level at 3 h Amp to 4 h Amp or 5 h Amp (long-term persister level). About 48 clones were grown from a single cell and treated with Amp (0.1 mg/mL), and the % survival was determined at different time points. r: Pearson’s correlation coefficient; ns: not significant.

### Cell density is not the primary factor for variation in persister number

Some bacterial persistence levels are controlled (in part) via quorum sensing^32^, which is a density-dependent response. Thus, before testing hypothesis 2, we wanted to know how much cell density could skew our results by testing the overall persister range with different cell densities using FTs. For FTs in Fig 2a, cells were harvested at OD 0.4-0.6 since the persister levels are remarkably similar at these ODs. We detected no correlation between cell density and % Survival in 15 FTs (7 Amp and 8 Apr) and a weak correlation in FT3 with Amp treatment (Fig. S1b *i*). Thus, to further determine how much cell density affects the persister levels, we tested persister levels from OD 0.3 to 0.7 in ∼0.1 OD intervals (cells are in log phase and cell density ranges from ∼2E07 to 2E08 CFU/mL) in a general population at 3 h Amp or Apr. No appreciable correlation was observed with either antibiotic (Fig. S1b *ii*).

### Cells are primed for persistence, not short-term tolerance

Short-term tolerance can mask persistence, and experimental evidence has shown that the phenotypes are distinct from each other^33^. Short-term tolerant cells are dividing and likely have different survival mechanisms than persisters. In the initial stage of antibiotic treatment, there are far more short-term tolerant cells than persister cells (Fig. 3 *i*). We did a similar FT with clones grown from a single cell and treated them with a lethal Amp concentration (0.1 mg/mL). % Survival was determined for 1 h and 3 h of treatment using persister assays^26, 27, 34^, and short-term tolerance at 1 h does not indicate the level of persistence at 3 h with lethal Amp; no correlation at 1 h vs 3 h population (r^2^ = 0.02). If the primed cells are advantageous for long-term survival, we expect the high persister populations observed at 3 h treatment to stay high with more prolonged antibiotic exposure. As expected, % Survival at 3 h highly correlates with % Survival at 4 h (r^2^ = 0.85) and 5 h (r^2^ = 0.78) Amp treated population (Fig. 3). Therefore, our results demonstrate that cells are primed for persistence and not for short-term tolerance.

### Primed cells have transient memory

Next, we determined whether high persister clones arise randomly due to noise in gene expression levels or in rare events where the gene expression level (memory) is passed down for several generations. We hypothesize that there is a transient memory at the transcriptomic level. The null hypothesis is that there is no memory and the variation range in persistence is random and solely due to noise. To test this, we divided and diluted the culture between 1:1 to 1:100 into separate microplates and allowed them to grow (Fig. 4a). If the null hypothesis is correct, there should be no correlation between divided cultures. However, persister levels were strongly correlated until the 1:20 dilution, and the memory is completely lost after a 1:100 dilution, supporting our transient memory hypothesis (Fig. 4b). We further confirmed the transient memory hypothesis with Apr (Fig. 4c). These results also add additional support that primed cells are not mutants, because 1:100 dilution led to no long-term (genetic) survival phenotype (in several different clones), as a resistance mutation would allow.

**Fig. 4.**
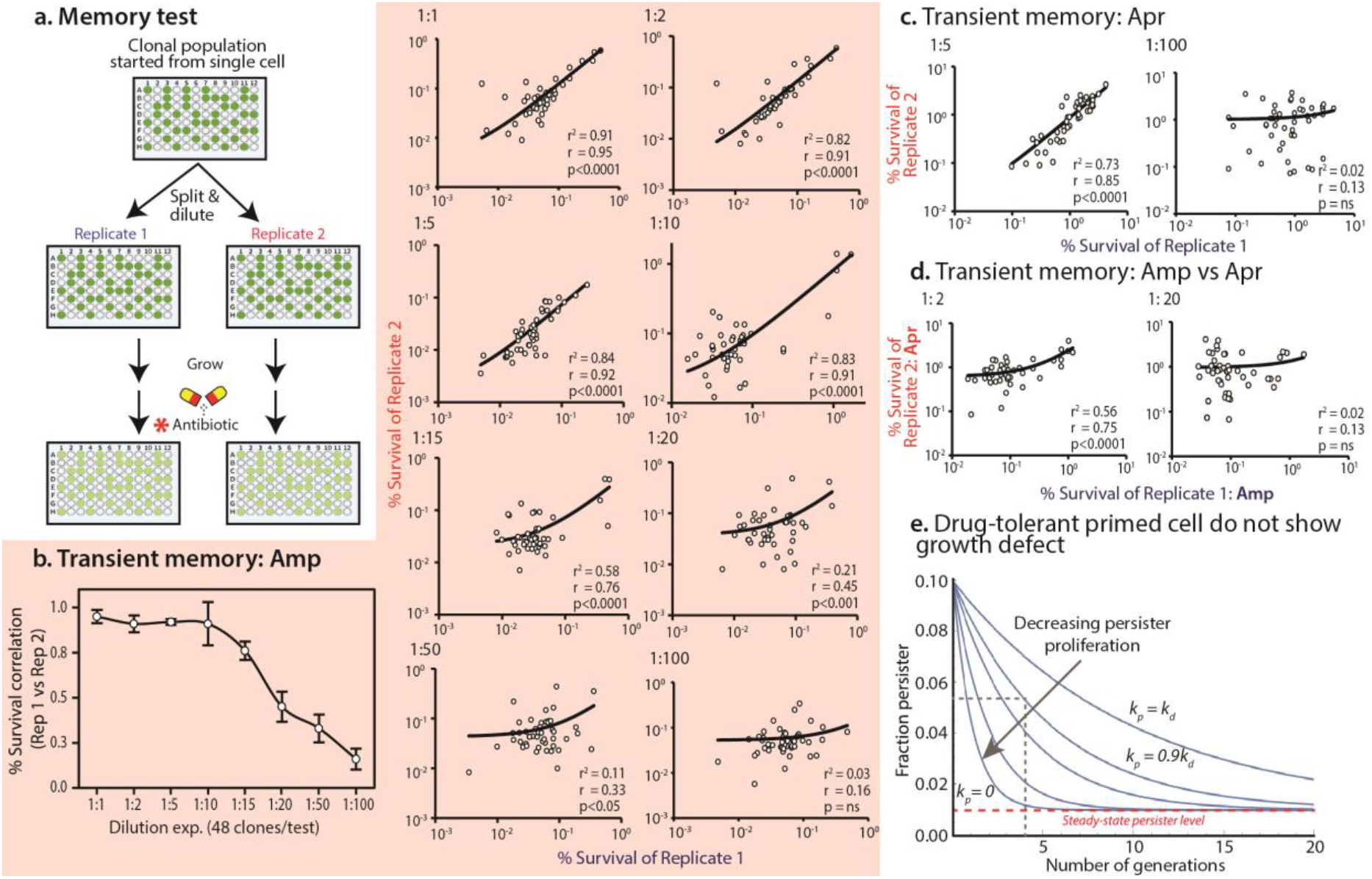
Primed cells have transient memory. **a**. Fluctuation test (FT) setup to check primed cell’s memory: cultures were diluted to 0.5 cells/well, grown to mid-log phase (∼1E+8 CFU/mL), each clone is divided and diluted into 2 replicates, grown to mid-log, and then tested for persistence. **b & c**. Primed cells have transient memory for several generations before 3 h Amp and 3 h Apr treatment. The persister levels correlate even with 1:20 dilutions between Replicate 1 and 2, suggesting a strong memory within the primed subpopulation. The memory is eventually lost with a 1:100 dilution. **d**. Primed cells show a weak memory when split and tested with Amp vs. Apr. *Antibiotic: Amp vs. Amp (replicate 1 & 2 both treated with Amp) or Apr vs. Apr (replicate 1 & 2 both treated with Apr) or Amp vs. Apr (replicates 1 & 2 treated with Amp and Apr, respectively). Axes are % Survival. r: Pearson’s correlation coefficient; ns: not significant. **e**. Plot representing the fraction of persisters 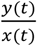 as predicted by the differential equation model (see Methods), assuming that initially 10% of cells where persisters and the steady-state persister level was *f*_*s*_ = 1%. The five lines are plotted assuming the persister proliferation rate, *k*_*p*_ = *k*_*d*_, 0.9*k*_*d*_, 0.8*k*_*d*_, 0.5*k*_*d*_, 0 where *k*_*d*_ = 1 and *k*_2_ = *k*_*p*_/10. The black dashed lines show the numbers of generations it takes for 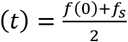, when *k* = 0.9*k*_*d*_.

We hypothesize that the same primed cells will allow higher persister levels when using antibiotics from classes that target distinct cellular processes, e.g. Amp and Apr. If both Amp and Apr primed cells use an akin mechanism, we expect a reasonable correlation between their persister numbers/well. If they do not correlate, diverse types of primed cells likely exist. To understand this, we did experiments similar to Fig. 4a, but we tested Replica 1 with Amp and Replica 2 with Apr. In this case, both replicas had a transient memory, and the memory was lost by 1:20 dilutions (Fig. 4d).

### Primed cells are not spontaneous persisters

Previous researchers proposed spontaneous persisters formation; persisters that are generated stochastically at a constant rate during exponential phase growth and switch to a dormant or a protected state (distinct slower growth rate than other cells, and this slowed growth rate is maintained for several generations and turn into persisters in the presence of stress)^5, 20, 35^. In addition, they proposed that during exponential growth, these phenotypic variants (e.g. persister formation) could also be induced by stress^5^. However, several research groups criticized the concept of spontaneous persister formation^36, 37^, questioning if it exists or is a proper terminology. In the original paper, where persistence was proposed in 1944, persisters were defined as non-dividing cells^38^. However, spontaneous persisters were defined as dividing cells^39^. Here we use the original definition of persisters; they do not divide. In addition, if persister formation is only induced by stress, all cells in the population should turn into persisters instead of a small percentage of the population. Also, induced persister formation (or sense-and-respond strategies) could be costly for the cells since it necessities constitutive expression of essential molecular machinery^40^.

On the contrary, primed cells (already prepared for stress in a population through heterogeneity) could provide a simple mechanism for adaptation to stresses they might or might not encounter. A key phenotype of spontaneous persisters is that they grow slowly. However, we did not observe any significant growth rate changes among the clones, and recent results demonstrate that persisters are not slow-growing before antibiotic treatment^41^. To further explore whether primed cells are reliant on slow growth, we constructed a simple mathematical model (explained in the Methods), where we plotted the fraction of persister cells 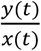 as a function of time for persister proliferation being 100%, 90%, 80%, 50%, and 0% of the proliferation rate of the drug-sensitive cells. We observed when persister cells do not proliferate (*k*_*p*_ = 0), their fraction rapidly dilutes back to the steady-state level in a few generations (Fig. 4e). These results show that a high correlation in persister maintenance, as seen in Fig 4b for several generations, requires persister proliferation. For example, one requires *k*_*p*_ = 0.9*k*_*d*_ for it to take roughly 4 generations for the fraction of persisters to fall to 50% of its initial levels, similar to the drop in the correlation between replicates to 0.5 in Fig. 4b. Our evidence clearly shows that primed cells are not persisters (non-dividing cells) before antibiotic treatment since they grow and maintain a transient memory.

## Discussion

Phenotypic heterogeneity is a fact of life and exists in both unicellular and multicellular organism^1, 2, 4^. This latent variation in phenotypic plasticity reveals in unfavorable environmental condition^4^. Noise in gene expression can drive this heterogeneity, and heterogeneity is hypothesized to be a key player that regulates bet-hedging strategies to endure harsh environmental fluctuation, such as bacterial persistence^2, 42^. Persisters have reduced efficacy to antibiotic treatment and are a key contributor to the rise in antibiotic resistance. Unraveling the underlying molecular mechanisms of heterogeneity is therefore crucial to comprehend bacterial persistence.

In this study, we used a powerful fluctuation test framework to infer transient cell states that arise via reversible and non-genetic mechanisms, recently employed in probing cancer persistence^13, 16, 17^, to find hidden features of bacterial persistence. Using the FTs, we tested the variation between clonal populations that originated from an identical clone and showed that a subset of the population, primed cells, have a “memory” that leads to high numbers of persisters. Our results demonstrate that under the same conditions, phenotypic heterogeneity often occurs in a range of ∼60-100 and ∼40-70 fold for Amp and Apr, respectively, despite (likely) being driven by a stochastic noise in gene expression. The relative consistency in primed cell numbers suggests that changes in specific genes result in primed cells, though we did not experimentally explore the mechanisms of primed cells in this work. Having high numbers of primed cells is a selective advantage that offers phenotypic plasticity to a bacterial population experiencing frequent harsh environmental stress. This heritable transient state might be favored in the course of evolution as a survival tactic compared to DNA mutation because it requires no long-term commitment. A recent paper demonstrated a transient cellular memory in *E. coli*, and that inheritance of non-genetic elements can help maintain cellular memory^43^. Thus, reducing variation among new cells for a few generations^43^. They found that some inherited elements, cell size, and the time required for cell division were maintained for nearly ten generations^43^.

We did not explore experimentally the regulatory mechanisms behind primed cells in this study, but we offered a mathematical model that supports and gives clues to the regulation of the transient memory. However, a few possible mechanisms that bacteria employ are already known. DNA methylation is a common method of epigenetic regulation in organisms, and many forms of methylation have been identified in bacteria^44-46^. Some are related to antibiotic resistance or persistence^47, 48^. For example, the deletion of *dam*, which is responsible for adenine methylation, can lead to lower persister numbers in pathogenic *E. coli* strains^49^. Another recent study showed multisite phosphorylation may regulate the phenotypic variability in a bacterial population. A gene encoding ppGpp Synthetase, *sasA*, is regulated by multisite phosphorylation of WalR and exhibits elevated levels of extrinsic noise in gene expression. Due to this noise, cells having elevated levels of *sasA* expression have increased short-term antibiotic tolerance^12^. Another possible mechanism could be ploidy. The Brynildsen group showed that ploidy or chromosome abundance produces phenotypic heterogeneity that affects persister numbers. Stationary-phase *E. coli* cells with two chromosomes had ∼40-fold more persisters than cells with one chromosome against fluoroquinolone antibiotic^50^.

In this study, we have not dealt with viable but nonculturable (VBNC) cells. Several studies showed VBNCs and persisters share similar phenotypes, and the major difference between them is after stress (e.g. antibiotics), persisters can grow on Petri plates while VBNCs can only grow in liquid. Currently, there is a furious debate if VBNCs are actually persisters, simply dying cells, or if VBNCs even exist^51-55^. A recent study called in to question many of the indicators that are currently used to classify bacterial cells as alive or dead^56^.

We tested if VBNCs could be resurrected using the 0.5 cell/well technique. The 0.5 cell/well is based on cell counts on Petri dishes (CFU/ml). Often, VBNCs are cited as being at equal concentrations (using a hemocytometer and microscope) as CFUs^57, 58^, but we did not observe this. We observed a ∼20% count difference (cell/mL) between hemocytometer and agar plate count, which is within the hemocytometer’s manufacture error rate. However, still if VBNCs are present and grow only in liquid, we expect 1 cell/well in a 96-wells plate when using CFU/mL from Petri dishes. Instead, we consistently get ∼48 wells had growth. We also consistently get growth in ∼24, ∼12, and ∼6 wells when we use 0.25, 0.125, and 0.0625 cells/well, respectively, with an r^2^ of 0.97, when plotting the wells with growth vs. cells/well based on CFU/ml. Supplementing media with pyruvate may resurrect VBNCs quicker^59, 60^. We tested the addition of pyruvate after antibiotic therapy with two carbon sources: glycerol and glucose. VBNCs did not resurrect; only ∼48 wells had growth.

Recognizing that the previous assays do not account for all the nuances of VBNCs, we tested whether VBNCs would alter our overall results using 2 cells/well (based on CFU/ml). Again, we see a high variation with Amp treatment, ∼78-fold (Fig. S1c.). This shows that if there are VBNCs, they are not skewing our results and that we still observe primed cells even with 2 cells in each well. It also demonstrates that the FT does not require 1 cell/well, which is an important finding because it is unrealistic to assume that every well we test will always have 1 cell when diluted to 0.5 cells/well.

Overall, our findings show that a subset of the population has a transient memory allowing them to prepare for antibiotic stress (we refer to them as ‘primed cells’). However, further exploration is needed to determine the regulatory mechanism(s) behind this cellular memory and the contribution of non-genetic factors in phenotypic heterogeneity.

## Methods

### Microbial strains and media

*Escherichia coli* DH5αZ1 having the p24KmNB82 plasmid was used in this study. We used this strain because DH5αZ1 was a derivative of *E. coli* K12 strain, and it has been used in our previous persistence studies^26, 27^. The cultures were grown in the defined media MMB+^26, 27^ with thiamine (10 μg/mL), 0.5% glycerol (or glucose), Km (25 μg/mL), and amino acids (40 μg/mL) or on Miller’s lysogeny broth (LB) agar plates +Km (25 μg/mL). For pyruvate assays, MMB+ media were supplemented with 2 mM sodium pyruvate. All cultures were incubated at 37°C and shaken at 300 rpm.

### Population-level variation check (Noise control: NC)

We used two different types of antibiotics, Ampicillin (Amp, target cell wall^30^) and Apramycin (Apr, target 30S ribosome^31^), for all fluctuation tests. We started with a mid-log phase *E. coli* culture having ∼1E+8 CFU/mL (∼OD 0.5), subsequently divided the culture into 48 well (48 replicate) of 96-Well Optical-Bottom Plate with Polymer Base (ThermoFisher) and treated them with Amp (0.1 mg/mL) or Apr (0.1 mg/mL) for 3 h at 300 rpm, 37°C in a FLUOstar Omega microplate reader. Persister percentage was calculated by comparing CFUs per milliliter (CFU/mL) before antibiotic treatment to CFU/mL after antibiotic treatment. Plates were incubated at 37ºC for 42-48 h, then scanned using a flatbed scanner. Custom scripts were used to identify and count bacterial colonies^61, 62^.

### Fluctuation test (FT) from a single cell level

We started with a log phase *E. coli* culture with ∼1E+8 CFU/mL (∼OD 0.5) and subsequently diluted the culture into 0.5 cells per well in a 96-Well Optical-Bottom Plate and grown in a FLUOstar Omega microplate reader at 37°C, 300 rpm. Each dilution experiment was done twice for each single-cell experiment to confirm the dilution, and cultures were only picked when 50-60% of the well had growth (48-55 wells out of 96 wells). The single cell was proliferated to mid-log phase ∼1E+8 CFU/mL (∼OD 0.5) and treated with Amp (0.1 mg/mL) or Apr (0.1 mg/mL) for 3 h, 300 rpm at 37°C. The persister percentage was calculated in the same manner described in the above section. The splitting and dilution test was done exactly as above, except when the single cell proliferated into mid-log phase, the cultures were separated and diluted into 2 plates with pre-warmed MMB+ media. Then, the cultures were grown to exponential phase, and persister assays were performed.

### Antibiotic resistance assay

After each fluctuation test, clones were diluted 1:100 in 1 mL LB media and were grown for 12 h. The cultures were then centrifuged for 3 min at 16,000 *x g*. After that, the supernatant was removed, and the pellet was streaked with and without antibiotic containing LB agar plate and incubated at 37°C.

### Minimal inhibitory concentration (MIC) test

Clones, including high persister clones from fluctuation tests, were diluted 1:100 in 1 mL LB media and grown for 12 h. First, 0.2 mL of culture was spread on an LB agar plate, then a MIC strip was placed on top of it and incubated at 37°C.

### Mathematical model

To understand primed cell proliferation rate, we considered a model of persister formation where drug-sensitive cells switch to a persister state with a rate *k*_1_, and persister cells revert back to the drug-sensitive state with a rate *k*_2_. We assumed the rate of cellular proliferation in the drug-sensitive and persisters states to be *k*_*d*_ and *k*_*p*_, respectively. In an expanding cell colony, the number of persister cells can be captured by the following system of ordinary differential equations,

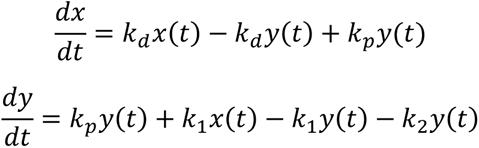

where *x*(*t*) and *y*(*t*) are the total number of cells and the number of persister cells, respectively, at time *t*. By setting,

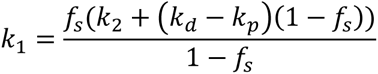

ensures that the steady-state persister fraction 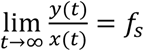. We considered a clone that initially had a high fraction of persister cells. We used the model to explore the relaxation of persister numbers back to steady-state levels, and how this time scale particularly depends on the persister proliferation rate *k*_*p*_. We then plotted the fraction of persister cells 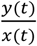 as a function of time for persister proliferation being 100%, 90%, 80%, 50% and 0% of the proliferation rate of the drug-sensitive cells.

### Cell counting

Initially, using the common protocol for a hemocytometer, microscopic counting, and live/dead dye, we saw about equal “VBNCs” per non-VBNCs in log and stationary phase, and with lethal antibiotics. This is consistent with the literature^57, 58^. However, we adjusted our counting protocol to match the manufacturer’s recommendation. As a result, we no longer see 2X the VBNCs compared to the CFU/ml; instead, the microscope count and Petri plate count only varied by ∼20 - 25% in log phase. We attempted to use several live-dead dyes to calculate viability after antibiotic treatment, but we found these assays to be quite unreliable, as recently cited in the literature^63-67^. So, we took the other approaches we described.

## Acknowledgements

This work is supported by the National Science Foundation award Numbers 1922542 and 1849206, and by a USDA National Institute of Food and Agriculture Hatch project grant number SD00H653-18/project accession no. 1015687. We would also like to thank KM Taufiqur Rahman for sharing his counting results using hemocytometer and petri plate assays.

## Author contributions

T.H. wrote the manuscript, performed the experiments, and ran data analysis. A.S. helped to formalize the research concept and design the experiments, as well as constructed the mathematical modeling. N.C.B. planned and directed the project. All authors contributed to discussing and editing the manuscript.

## Supplement Figure

**Supplement Fig. S1.**
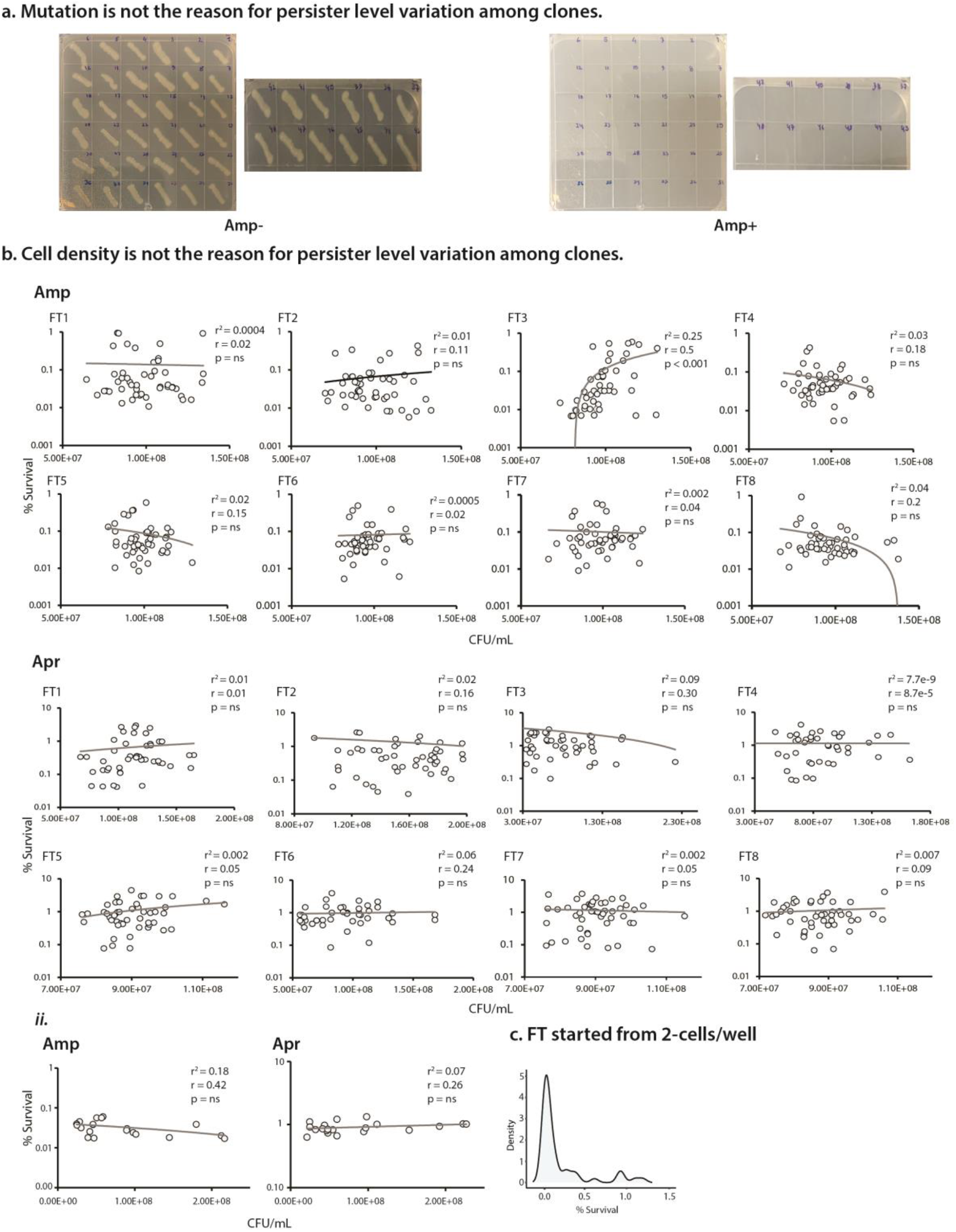
**a**. Streaking plate shows mutation is not the reason for persister level variability among clones. **b**. Cell density is not the reason for persister level variation among clones. ***i***. Each FT tests treated with Amp or Apr show no correlation between CFU/mL vs. % Survival. ***ii***. General populations treated with Amp or Apr show no correlation between CFU/mL vs. % Survival. **c**. FT started with 2-cells/well, and showed ∼78-fold variation among clones.

